# Bacterial Community Composition and Diversity along the Southern Coastlines of the Atlantic Ocean in Cape Town, South Africa

**DOI:** 10.1101/2020.02.27.969030

**Authors:** Ola A. Olapade

## Abstract

The spatial distribution and diversity within bacterioplankton assemblages in four coastal sites along the southern points of the Atlantic Ocean were examined using the Illumina high-throughput that targets 16S rRNA genes to examine indigenous bacterial assemblages in the littoral zones along the coast of the ocean. Results of the study showed very similar bacterial representation between the coastal sites with majority of the sequences affiliated with *Proteobacteria* (52 to 59%), *Bacteriodetes* (21 to 31%) followed by *Actinobacteria* (3 to 9.5%) and *Planctomycetes* (2.1 to 4.5%). The bacterioplankton assemblages at each site examined were quite diverse, with members of the *Gammaproteobacteria* found as the most abundant bacterial class among the four sites. However, clear differences were observed among the sites at the order level, with the *Chromatiales* the more dominant in the closer CPTI sites, while clades belonging to the *Flavobacteriales* and *Rhodobacterales* were more prevalent in the two CPTA sites. While the results of UPGMA clustering and principle coordinate (PCoA) revealed two spatially separate clusters among sites, canonical correspondence (CCA) analysis indicated that environmental variables such as temperature, pH and conductivity were probably the major influencers of bacterial occurrences at the coastal sites.

## Introduction

The bacterioplankton assemblages in oceans have been described as comprising one of the largest and active microbial assemblages in the biosphere (e.g., Whitman et al 1998), where they actively partake in the biogeochemical influxes and cycling of various nutrients and organic compounds (Azam and Malfatti 2007, Falkowski et al 2008, Zehr and Kudela 2011). Despite the well documented knowledge regarding the occurrence and productivity of bacterial assemblages in oceans, yet several questions still remain unanswered about their structural composition, spatial distributions and diversity (e.g., Salazar and Sunagawa 2017). For instance, contrasting findings have been previously reported regarding bacterial diversity and biogeographic distributions in marine systems, especially in tropical oceans (e.g., Pommier et al 2007, Fuhrman et al. 2008, Milici et al. 2016). Milici et al. (2016) reported a clear biogeographic pattern with a double inverted latitudinal gradient, with higher diversity in planktonic bacteria population in mid-latitudinal regions, and decreasing towards the equator in the Atlantic Ocean in their study. In contrast, Fuhrman et al. (2008) found a negative correlation between species richness and latitude both in the Northern and Southern hemisphere. Additionally, interesting observations regarding microbial provincialism and their discrete distributions in various marine habitats have been previously documented (e.g., Brown et al. 2009, Jeffries et al. 2015).

In order to better understand the structural compositions and diversity within bacterial assemblages in marine systems, this study was conducted on several coastal sites along the southern points of the Atlantic Ocean in Cape Town, South Africa by examining indigenous populations in the bacterioplankton communities using Illumina high-throughput sequencing approach that targets the 16S ribosomal RNA genes. The four sites that were selected for the study are located along the coastline within the metropolitan city of Cape Town, between the Cape of Good Hope and the Cape of Agulhas two touristy landmarks that are prone to various anthropogenic influences. The main aim of the study was to examine the taxonomic profiles of the microbial assemblages indigenous to these coastal marine sites as well as determine the influences of various environmental factors, such as temperature, pH and dissolved oxygen concentrations on the structural composition and diversity within the assemblages at the four sites. The Atlantic Ocean is considered the second largest of the world’s five oceans, second only to the Pacific Ocean, with a body of water located between Africa, Europe, the Arctic, America and the Southern Ocean. The southern parts of the Atlantic Ocean where this study was conducted is located around the metropolitan area of the city of Cape Town in South Africa, between the Cape of Good Hope (∼34 ° 21’24.63” S, 18 ° 28’ 26.36” E) and Cape Agulhas (∼34 ° 49’ 59.6” S, 20 °00’ 0” E), with mostly rocky headlands of coastlines in between these two landmarks that are about 90 miles apart (International Hydrographic Organization, 2002).

## Materials and Methods

### Sample Collection and Measurement of Environmental Variables

Samples were collected from the surface waters along the rocky headland coasts of the Atlantic Ocean in Cape Town, South Africa in September 2019. Specifically, water samples were collected from approximately 1 to 5 m depth into sterile falcon tubes from four separate sites along the ocean front close to the metropolitan city of Cape Town in South Africa (Figure 1). Collected samples were filtered through 0.2-µm pore-size polyethersulfone membrane filters and then stored frozen until nucleic acid extraction was performed. While on site, the water chemistry properties were also measured (see Table 1) with probes for temperature, conductivity, pH, dissolved oxygen and oxidation-reduction potential using the YSI model 556 MPS multi-probe system (YSI Incorporated, USA).

**Table 1:** Environmental variables measured at the study sites.

| Site | Latitude | Longitude | Temperature<br>°C | Conductivity<br>mS/cm | DO<br>% | pH | ORP |
| --- | --- | --- | --- | --- | --- | --- | --- |
| CPTA1 | 34° 21' 25." S | 18° 28' 26" E | 14.8 | 48.45 | 127.6 | 8.92 | -26.5 |
| CPTA2 | 34° 21' 24." S | 18° 28' 36" E | 14.76 | 48.5 | 111 | 8.8 | -16.5 |
| CPTI1 | 34° 13' 36. " S | 18° 28' 7." E | 14.93 | 48.93 | 389.2 | 9.14 | -23.6 |
| CPTI2 | 34° 49' 59. " S | 18° 28' 7." E | 14.89 | 48.45 | 263.3 | 9.07 | -20.5 |

**Figure 1.**
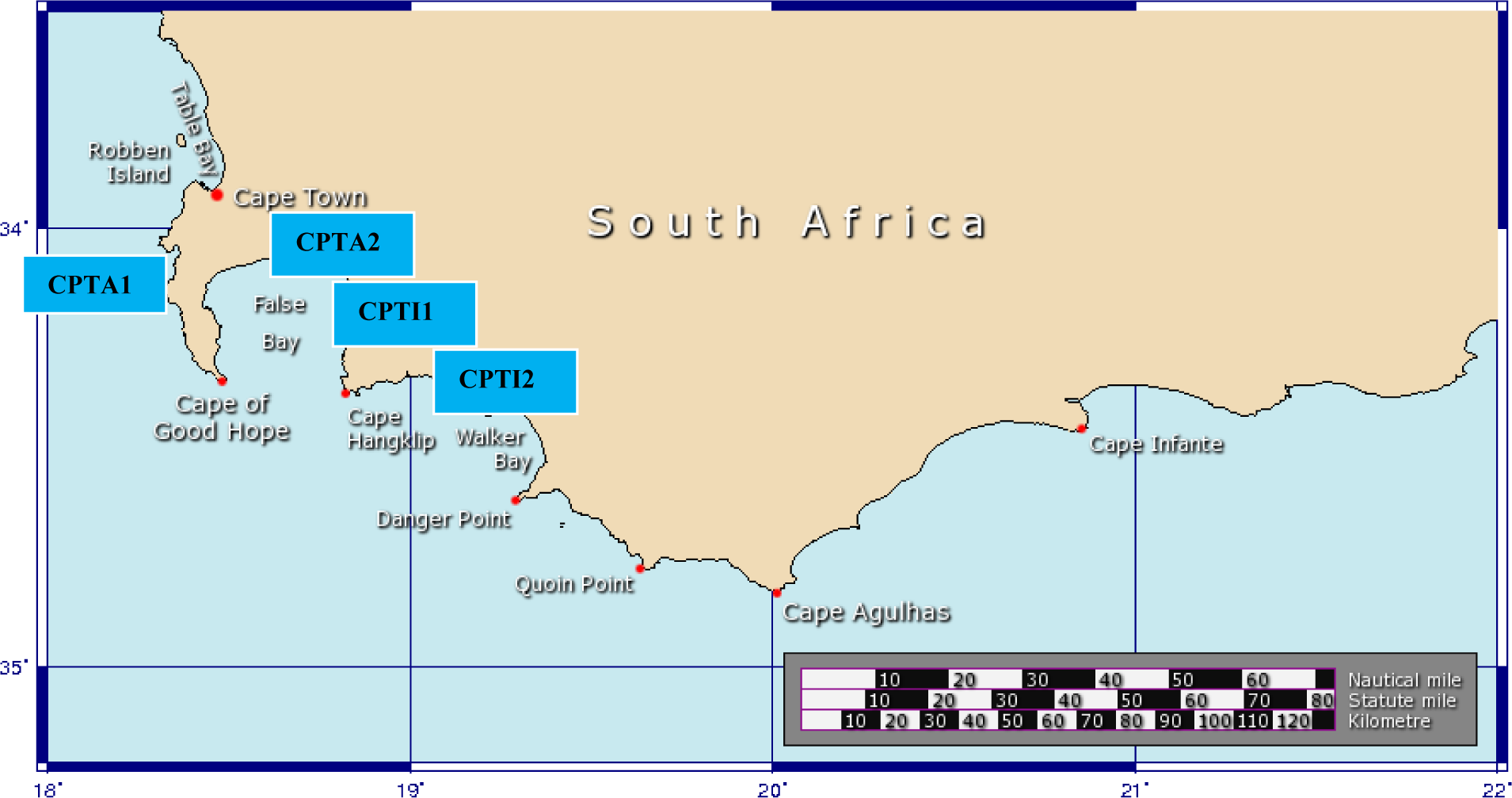
Map of study sites along the coastlines of the Atlantic and Indian Oceans in Cape Town, South Africa

### DNA Extraction and 16S rRNA Gene Pyrosequencing

Community DNA was extracted from the filters using FastDNA SPIN Extraction kit (MP Biomedicals, Solon, OH, USA) and eluted in 50 uL of sterile deionized water according to the vendor’s instructions. Determination of DNA quantity was then carried out with a NanoDrop Spectrophotometer (NanoDrop 2000, Thermo Scientific, Delaware, USA). The quality of extracted DNA was further assessed by amplifying with the 16S rRNA universal primer sets, 27F (5’ AGA GTT GTA TCM TGG CTC AG 3’) and 1492R (5’GGT TAC CTT GTT ACG ACT T3’) as previously described in Olapade (2013) and (2015).

The Illumina’s 16S metagenomic sequencing library preparation protocol was used in generating amplicon libraries using universal primer pairs that consisted of an Illumina-specific overhang sequence and locus-specific sequence: 92wF_Illum: 5’-TCGTCGGCAGCGTCAGA TGTGTATAAGAGACAGAAACTYAAAKGAATTGRCGG and 1392R_Illum: 5’GTCTCGT GGGCTCGGAGATGTGTATAAGAGACAGACGGGCGGTGTGTRC. The pair of primers targets the V6-V8 hypervariable regions of 16S rRNA genes of all microbial groups (Jefferies et al. 2015).

### Quality Trimming and Filtering of Low-Quality Sequences

The raw pyrosequencing data was processes and analyzed using the open-source software program, Mothur (Mothur v. 1.36.1; http://www.mothur.org) as previously described (Schloss et al. 2009). Barcode and the fusion primers are trimmed before any of the bioinformatics commences. Sequences reads without a barcode or a primer region are dropped and not considered for further analysis. Low quality sequences i.e. those less than 300 base pairs as well as those with less than average quality score (value of 25 or less) are filtered out and deleted (Zhang et al. 2012). Operational taxonomic units (OTUs) were constructed by comparing them to close relatives via global pairwise alignment (Altschul et al. 1997) to determine their close relatives using the BLASTN system. Chimeras were detected in the sequences that were later omitted for further analysis by using the UCHIME program (Edgar et al. 2011).

### Statistical and Diversity Analysis

The sequences were clustered into OTUs after setting 97% distance limit or cutoff similarity value (Tindall et al. 2009; Edgar et al. 2011). The OTUs were analyzed for species richness, Shannon Index, Simpson’s (Reciprocal) Index of diversity, species evenness, ACE richness estimate and Chao-1 richness indicator (Chao 1984; 1987; Chao and Lee 1992; Schloss and Handelsman 2006). In order to determine whether total diversity was covered by the numbers of sequences screened, Good’s Library Coverage values were calculated using the equation: C = 1 – (n/N) X 100, where n is the number of unique OTUs and N is the total number of clones examined as previously described (Good 1953; Kemp and Aller 2004). Alpha, beta and gamma diversity calculations were carried out according to Whittaker (1972); in addition to rarefaction analysis that was performed to also determine the diversity of the clone libraries using the freeware program by CHUNLAB Bioinformatics Made Easy (CLcommunity version 3.30). Taxon exclusive (XOR) analysis was also carried out based on the taxonomic assignment of sequencing read on the sets of clones obtained from the microbial assemblages to reveal those sequences present in one library but absent in the others as described by Li and Godzik (2006). The UPGMA Fast UniFrac analysis was used to cluster the sequenced microbial communities based on phylogenetic relationship and abundance in order to generate a dendogram (Hamady et al. 2010), while the multi-dimensional UniFrac distance matrixes were then converted into vectors using the Principal coordinate analysis (PCoA) as described by Jolliffe (1989). Additionally, canonical correspondent analysis (CCA) was also used to analyze and examine which of the bacterial assemblages corresponds to the independent environmental variables that were measured at the study sites according to Ter Braak and Verdonschot (1995).

The Student t-tests and ANOVA analyses were performed to analyze the differences in water chemistry characteristics among the four coastal sites examined with SPSS for windows (version 23, SPSS Inc., Chicago, IL). Post-hoc tests were also carried out for pair-wise comparison using Tukey HSD test. All data sets were log-transformed before performing statistical analyses. Statistical significance was set at *p* ≤ 0.05.

## Results

### Environmental Variables

The environmental variables measured at the four coastal sites along the ocean front included temperature, pH, dissolved oxygen (DO), conductivity and oxidation-reduction potential (ORP). Most of these variables were quite similar among the studied sites, with the exception of DO that was relatively higher in the two eastern sites (CPTI1 and CPTI2) closest to the Indian Ocean. Specifically, water temperature among the four coastal sites ranged from 14.76 to 14.89 °C and slightly higher in the two easterly located sites, while pH was also in the range of 8.8 to 9.14 between the western and eastern sites, respectively (Table 1).

### Community Composition and Diversity Analysis

Based on the 16S ribosomal RNA gene sequencing, the relative abundance of bacterial taxa was determined at different taxonomic levels, and majority of the bacterial sequences (∼99%) were ascribed to 29 different known bacterial phyla. Out of these 29 phyla, bacterial members belonging to the *Proteobacteria, Bacteroidetes, Actinobacteria, Planctomycetes* and *Verrucomicrobia* accounted for more than 90% of total community compositions among the sequences obtained from the four coastal ocean sites (Figure 2A). Members of the *Proteobacteria* were the most numerically dominant phyla in the four sites, accounting for between 52 to 59% of total bacterial sequences. The *Bacteroidetes* were in close second with between 21 to 31% sequence representations, followed by members of the *Actinobacteria* and *Planctomycetes* (between 3 to 9.5% and 2.1 to 4.5%, respectively). The next 25 bacterial phyla (including the members of the *Acidobacteria, Firmicutes, Tenericutes, Rhodothermaeota, Cyanobacteria, Deinoccocus-Thermus, Gemmatimonadetes*, among others) represented less than 10% of the total bacterial sequence abundance. Among the *Proteobacteria*, members of the *Gammaproteobacteria* were the predominant class, representing between 32 to 34%, followed closely by the *Alphaproteobacteria* with 13 to 14% representation among the four studied sites (Figure 2B). The *Chromatiales* were the dominant groups among the members of the *Gammaproteobacteria* at the order level, followed closely by the *Flavobacteriales* and the *Rhodobacterales*, both members of the *Flavobacteria* and *Alphaproteobacteria* classes, respectively (Figure 2B).

**Figure 2A.**
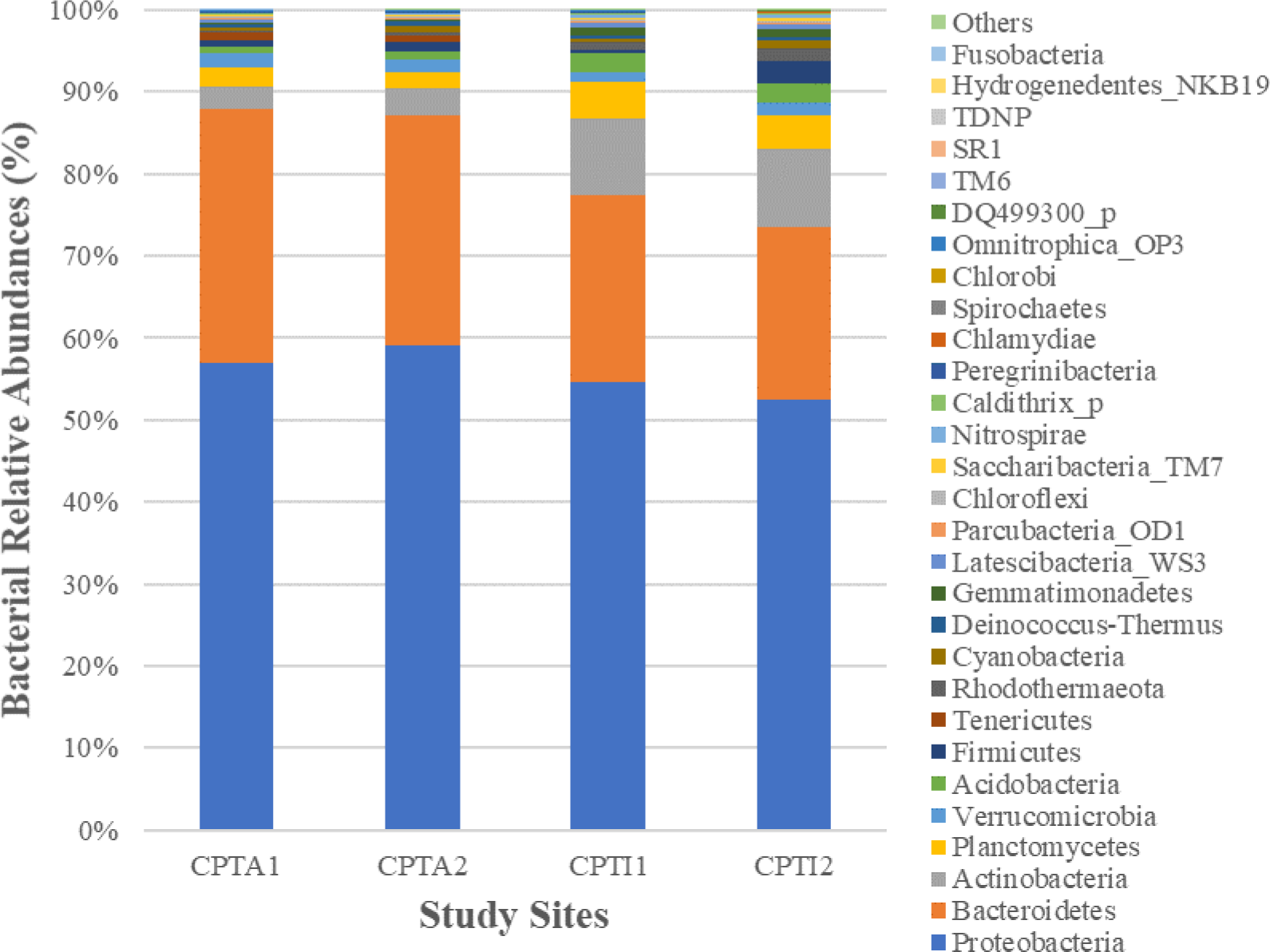
Relative abundances of bacterial phylogenetic taxa at the phylum level

**Figure 2B.**
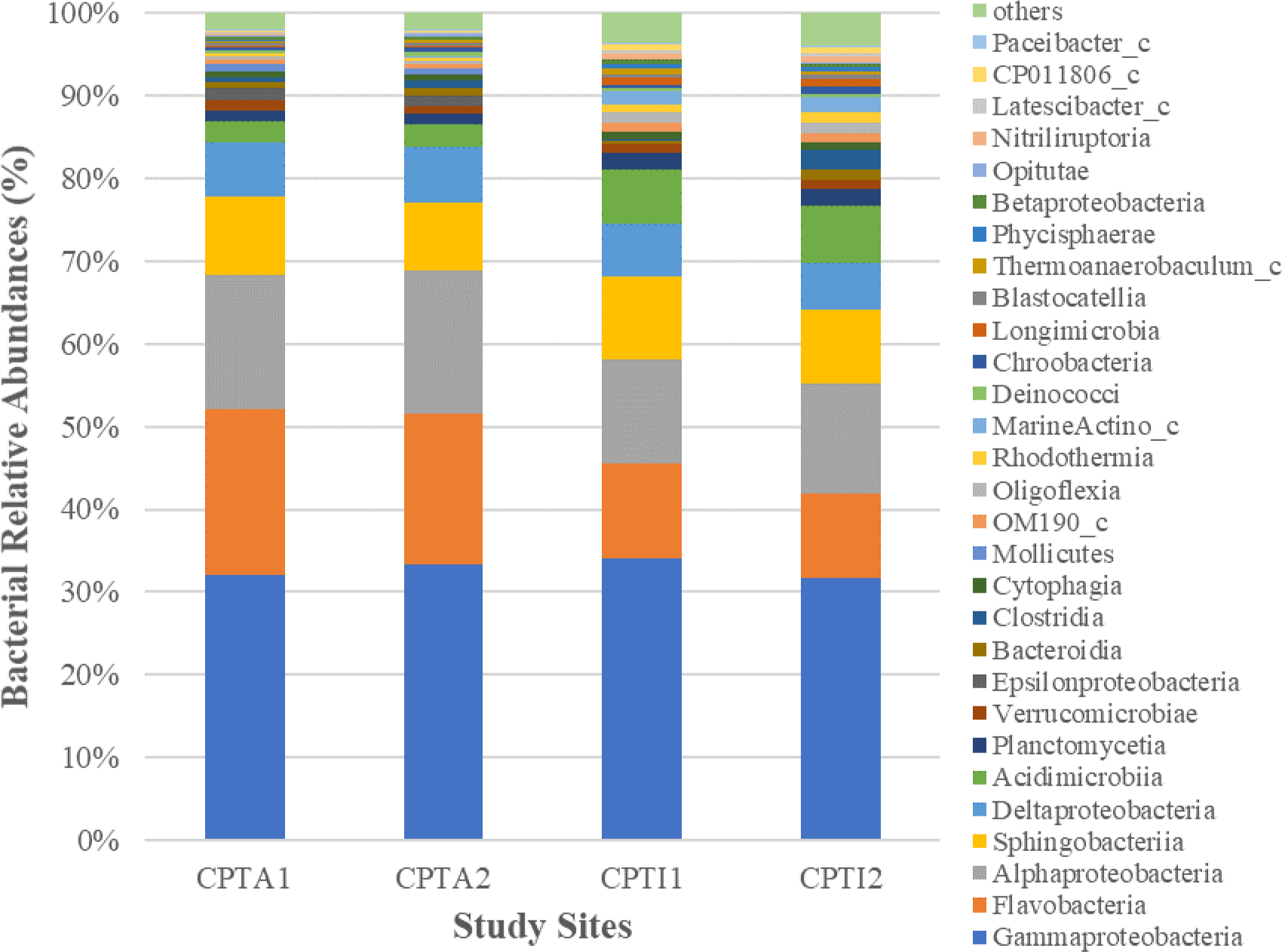
Relative abundances of bacterial phylogenetic taxa at the class level

**Figure 2C.**
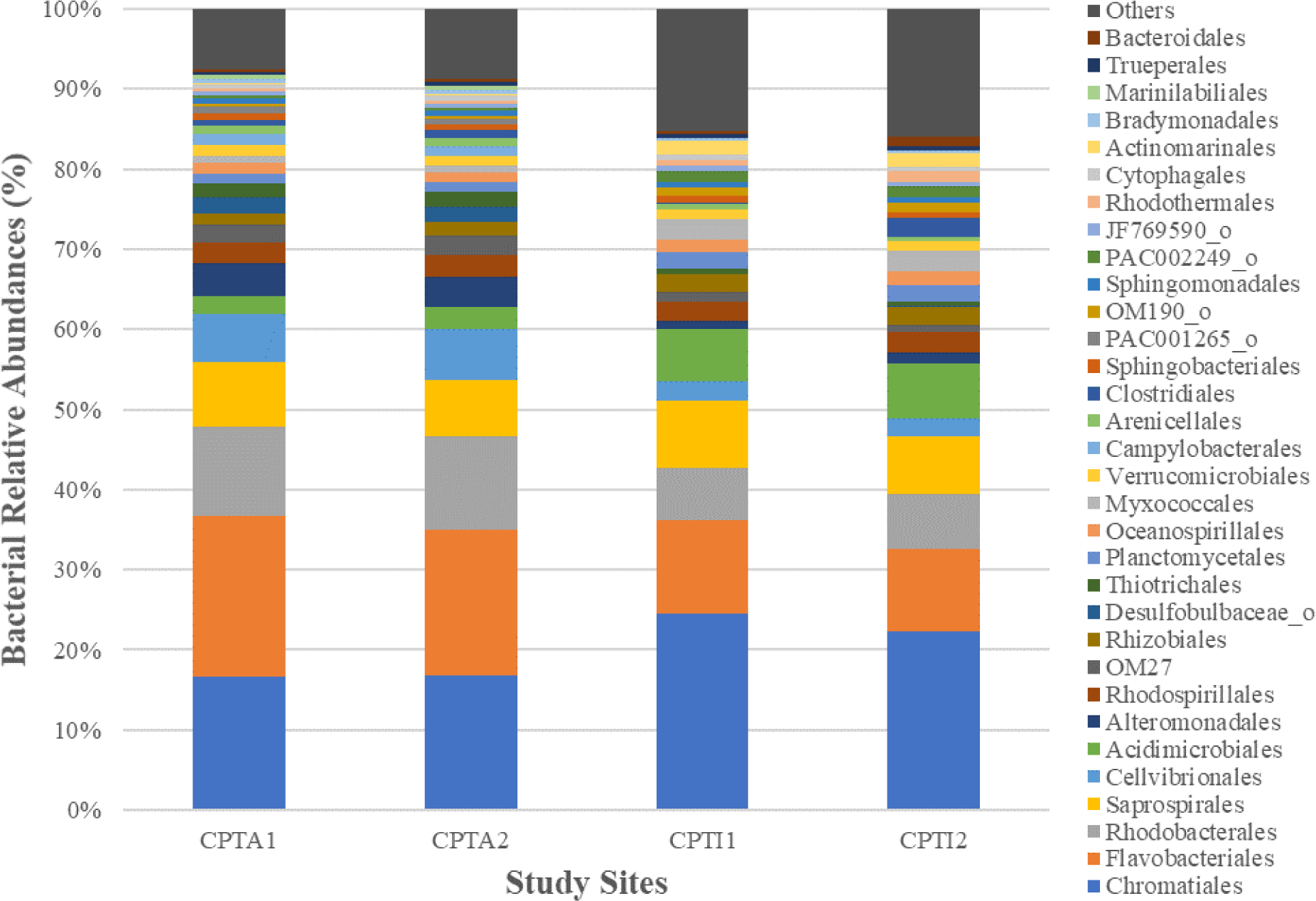
Relative abundances of bacterial phylogenetic taxa at the order level

The Good Library Coverage analysis revealed that majority of the bacterial sequences were covered among the sites (Table 2). This result is also corroborated by the rarefaction curves that showed sufficient coverage in the numbers of the different bacterial phyla within the four assemblages at the sites (Figure 3). Diversity measures such as the Shannon diversity index showed that bacterial diversity was comparatively higher in the two easterly located sites (CPTI1 and CPTI2) than in the two western sites examined. Results of bacterial richness and species diversity based on ACE and Chao1 while not showing a distinct delineation among the four coastal sites, however revealed that the CPTA1 site had the highest diversity compared to the three other sites. This result is further validated by the results of the taxon exclusive analysis of the bacterial assemblages at the phylum level that showed disparate differences in bacterial diversity among the sites (Table 3). The bacterial assemblages found within the CPTA1 coastal site comprised of several bacterial phyla that were totally absent in the three other sites including the *Armatimonadetes, TM6, Lentisphaerae, BRC1, TDNP, SR1, Elusimicrobia, Fibrobacteres and WS6*. While the CPTA2 site comprised of *Deferribacteres* that were absent in the other sites, and phyla belonging to *Aminicenantes, GN04* and *Synergistetes* were exclusively found in CPTI1 and CPTI2, respectively.

**Table 2:**
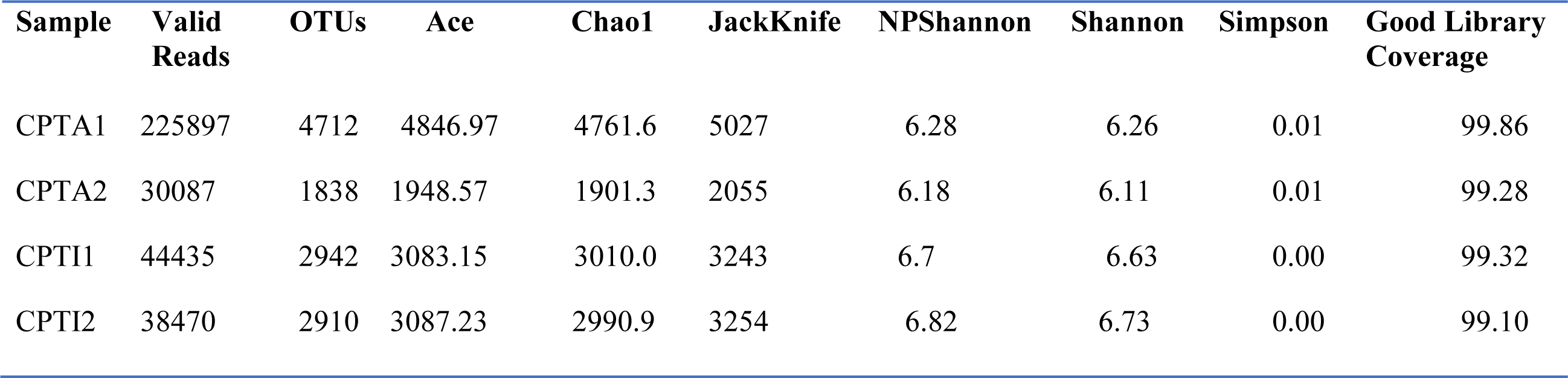
Community diversity analysis of the 16S ribosomal RNA gene sequences from the Bacterioplankton of the Atlantic and Indian Oceans in Cape Town, South Africa

| Sample | Valid Reads | OTUs | Ace | Chao1 | JackKnife | NPShannon | Shannon | Simpson | Good Library Coverage |
| --- | --- | --- | --- | --- | --- | --- | --- | --- | --- |
| CPTA1 | 225897 | 4712 | 4846.97 | 4761.6 | 5027 | 6.28 | 6.26 | 0.01 | 99.86 |
| CPTA2 | 30087 | 1838 | 1948.57 | 1901.3 | 2055 | 6.18 | 6.11 | 0.01 | 99.28 |
| CPTI1 | 44435 | 2942 | 3083.15 | 3010.0 | 3243 | 6.7 | 6.63 | 0.00 | 99.32 |
| CPTI2 | 38470 | 2910 | 3087.23 | 2990.9 | 3254 | 6.82 | 6.73 | 0.00 | 99.10 |

**Table 3:** Result of taxon exclusive analysis at the phylum level to detect taxa that are present in one bacterioplankton assemblage but absent in the others based on 16S rRNA gene sequences/clones

| Taxonomic Group<br>by<br>Phylum | Number of<br>Sequences/Clones |  |  |  |
| --- | --- | --- | --- | --- |
|  | CPTA1 | CPTA2 | CPTI1 | CPTI2 |
| *Armatimonadetes* | 3 | 0 | 0 | 0 |
| *TM6* | 56 | 0 | 2 | 3 |
| *Lentisphaerae* | 6 | 0 | 17 | 7 |
| *BRC1* | 15 | 0 | 4 | 2 |
| *TDNP* | 40 | 0 | 0 | 0 |
| *SR1* | 47 | 0 | 0 | 0 |
| *Elusimicrobia* | 10 | 0 | 4 | 3 |
| *Fibrobacteres* | 13 | 0 | 0 | 0 |
| *WS6* | 5 | 0 | 0 | 0 |
| *Deferribacteres* | 0 | 3 | 0 | 0 |
| *GN04* | 0 | 0 | 4 | 0 |
| *Aminicenantes_OP8* | 0 | 0 | 1 | 0 |
| *Synergistetes* | 0 | 0 | 0 | 2 |
| *Bacteria_uc* | 2 | 0 | 0 | 0 |

**Figure 3.**
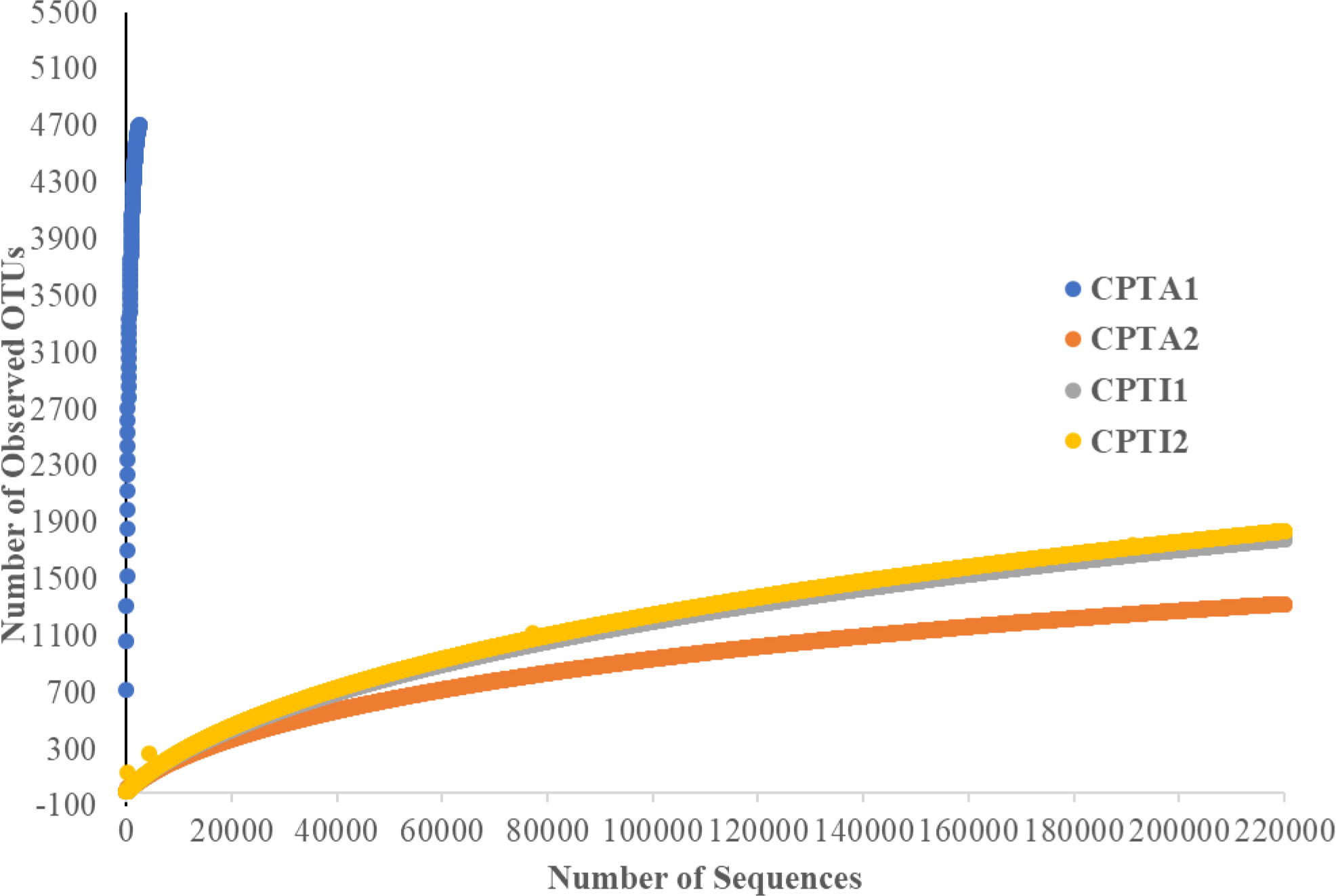
Rarefaction curves of OTUs based on 16S rRNA sequences from the bacterial assemblages from the study sites

Hierarchical clustering based on the Fast UniFrac distance matrix revealed that the bacterial sequences obtained from the two bacterioplankton assemblages in the western Atlantic Ocean sites were more similar, but were a bit distant from those in the easterly located assemblages (Figure 4). The PCoA that was also carried out to further explain the variations in bacterial community compositions between the four coastal sites also corroborated the results of the UPGMA clustering. Three axes were extracted that together explained 90.1% of the observed variance and showed that the bacterial assemblages within the two western sites (CPTA1 and CPTA2) clustered along the PC1 axis, while those from the eastern sites of the ocean (CPTI1 and CPTI2) clustered around the PC2 axis (Figure 5).

**Figure 4.**
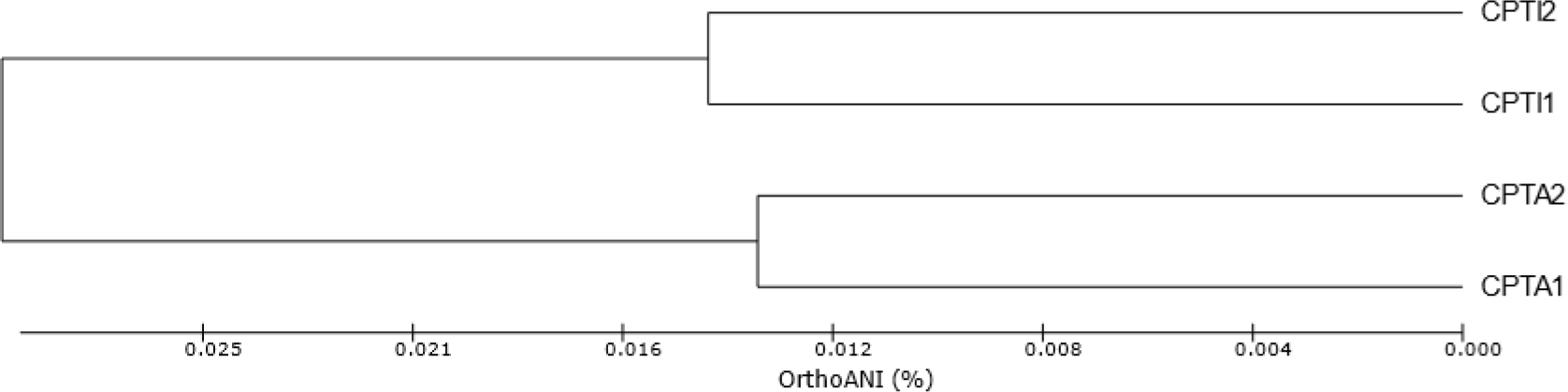
UPGMA (Unweighted pair group method with arithmethic mean) dendogram showing the clustering of bacterial assemblages from the study sites

**Figure 5.**
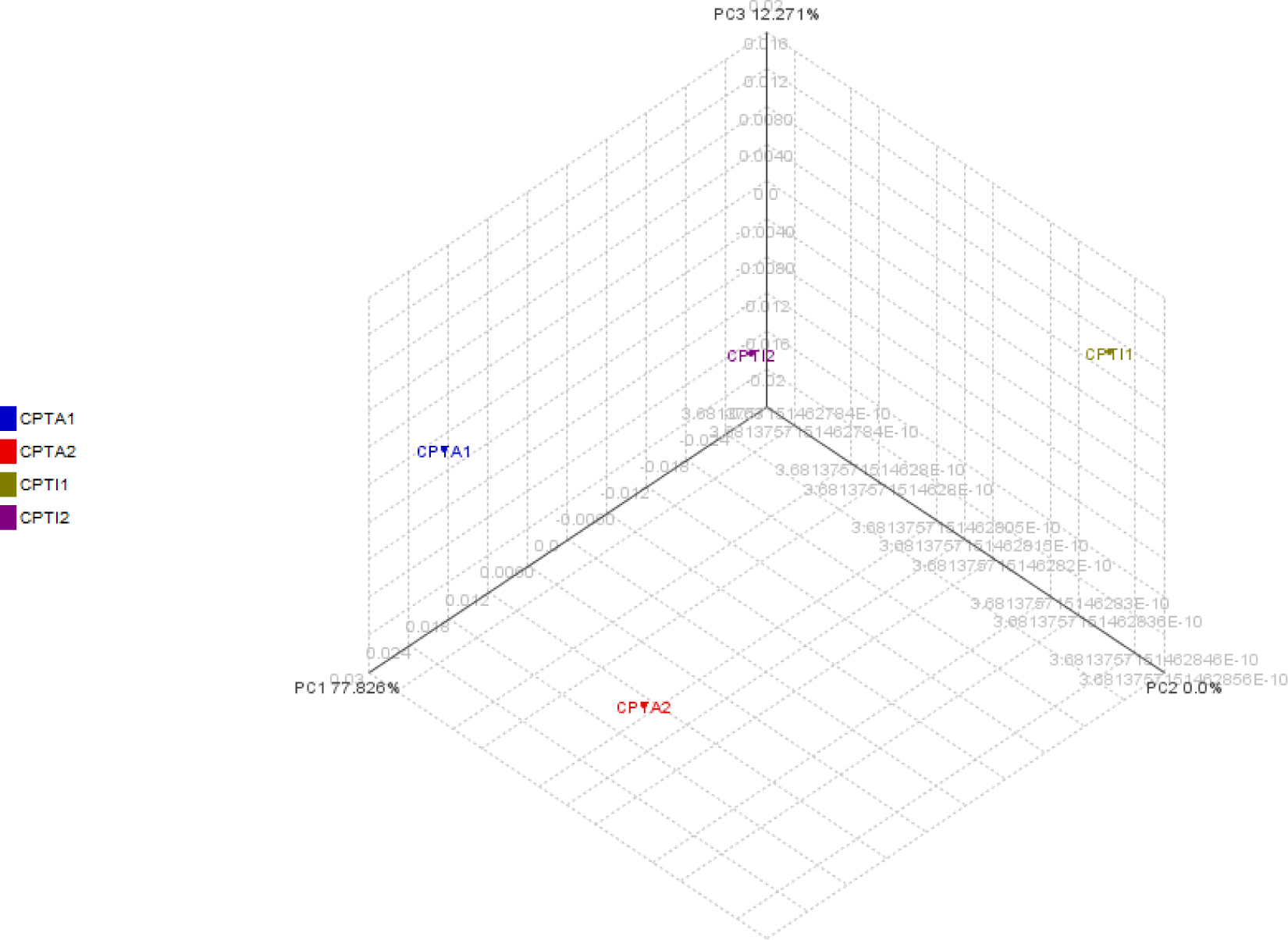
Three-dimensional principal coordinate analysis (PCoA) based on the Unifrac distance matrix of the bacterial assemblages for normalized OUT abundances within the study sites

CCA was carried out to better understand bacterial distribution patterns along the coastal sites, especially regarding the spatial occurrences of the various environmental factors that were measured. Therefore, temperature, pH, conductivity, DO and ORP were included in the CCA analysis. The environmental variables in the two CCA axes (i.e. CCA1 and CCA2) together explained more than 98.46% of total variations in the bacterial abundance distribution (Figure 6). Temperature, pH, conductivity and DO (p < 0.01) all contributed significantly to the total variance and were closely associated with the first and second CCA axes.

**Figure 6.**
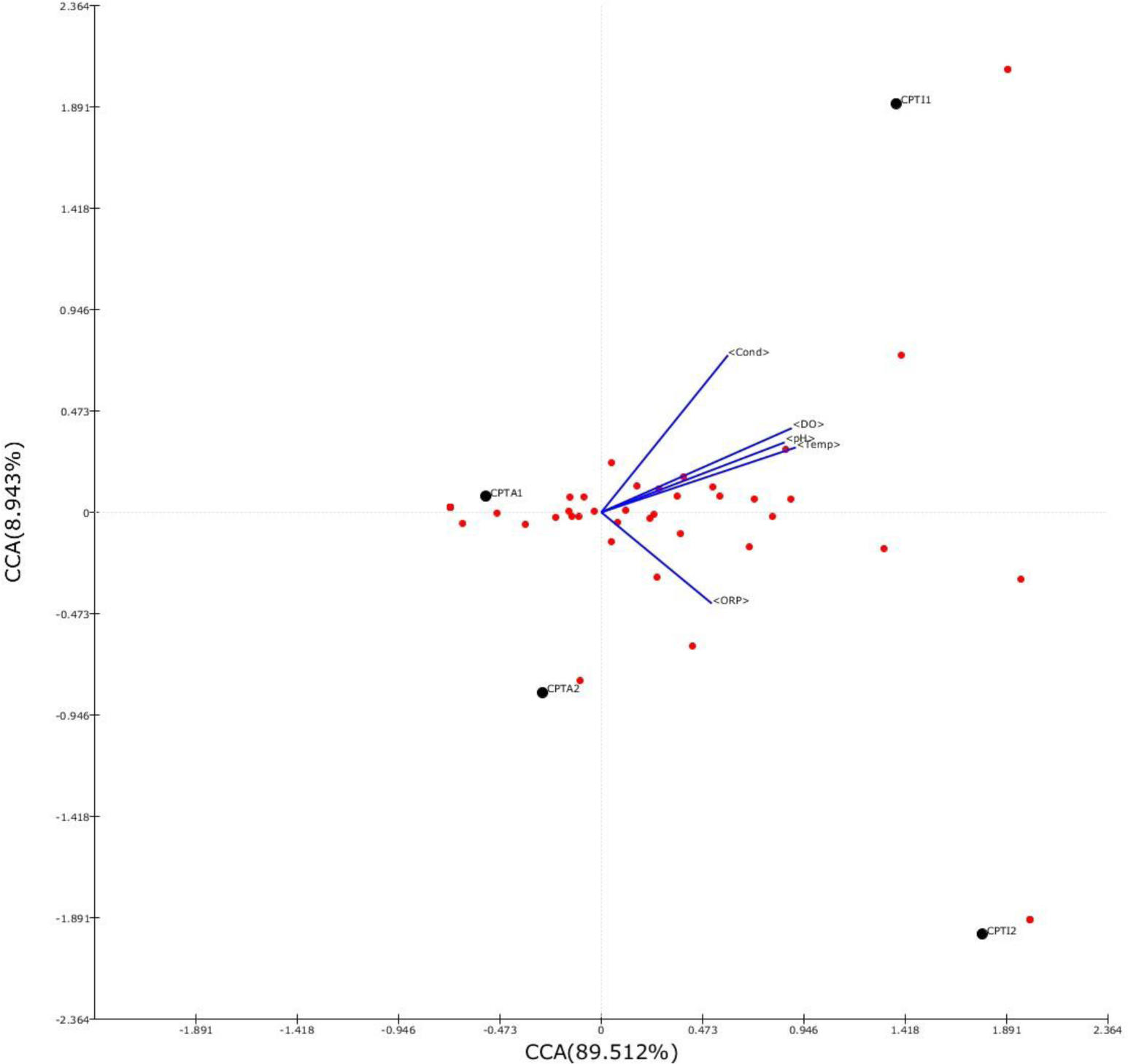
Canonical Correspondent analysis (CCA) of the bacterioplankton assemblages shown relationships with environmental variables within the study sites examined

## Discussion

In this study, the 16S rRNA gene sequences obtained from 4 different coastal locations along the southernmost parts of the Atlantic Ocean in Cape Town, South Africa were analyzed in order to characterize the bacterial community structures in response to potential changes in environmental variables between these spatially different coastal marine sites. The 16S rRNA occurrences diverse phylogenetic groups within the assemblages revealed close similarity in the dominant taxa among the four coastal sites examined. Bacterial members at each of the sites were mostly dominated by the *Gammaproteobacteria* class followed by the *Flavobacteria*. The numerical dominance of members of the *Proteobacteria* and *Bacteroidetes* as observed in this study is consistent with previous studies that have also reported the high occurrences of these phyla in marine systems (e.g., Brown et al. 2009, Seo et al. 2017, Wang et al. 2018, Wu et al. 2019).

The relatively high occurrences of members of the *Gammaproteobacteria, Alphaproteobacteria* and *Flavobacteria* among the sequences in the four coastal sites examined in this study is fairly consistent with those reported for oceanic waters by previous studies (Kirchman 2002, Rappe and Giovannoni 2003, Schmidt et al. 1991, Raes et al. 2017, Wang et al. 2018, Wu et al. 2019). These bacterial groups are known to be major constituents of microbial assemblages in various marine systems (e.g., Kirchman 2002, Rappe and Giovannoni 2003), especially in coastal environments because of their propensity for the high availability of enhanced dissolved organic matter that are copiously produced by photosynthetically active autotrophs in this euphotic area of the ocean. Previous studies have revealed significant numbers of these phyla strong correlations with dissolved organic matters associated by phytoplankton (Calson et al. 2009) as well as other environmental variables, such as phosphate concentration (Morris et al. 2010, Seo et al.2017) as well as salinity and temperature (Milici et al. 2016, Wu et al. 2019) in coastal marine waters. Milici et al. (2016) particularly found bacterial diversity to change drastically with changing water temperature in the Atlantic Ocean, with the highest at between 15 and 20°C, but decreased significantly when water temperature was above 20°C. The results from this study seems to validate their observations, given that the temperature of the four sites examined here were on average around 15°C and most of the diversity measures, especially as indicated by both ACE and Chao1 showed species richness to be high in all the sites examined.

The CCA results of this study showed that combinations of the major environmental variables probably explain the distribution of the different bacterial phyla among the coastal sites examined. Although other variables such as inorganic nutrients, including phosphate and nitrate concentrations were not included in the analysis conducted in this study, however other previous studies have shown strong correlations between their concentrations and the spatial distributions and diversity of bacterial assemblages in marine waters (e.g., Raes et al. 2017, Seo et al. 2017). Raes et al. (2017) found strong correlations between total dissolved inorganic nitrogen, chlorophyll a, phytoplankton community structure and primary productivity with bacterial richness in their study on the surface waters of the eastern Indian Ocean. While Seo et al. (2017) reported significant influences of both phosphate and dissolved oxygen concentrations on the bacterial community compositions found in different stations, mostly in the coastal waters, in the South Sea of Korea.

In general, the results from this study on the occurrences, distribution and diversity of bacterial populations within coastal bacterioplankton assemblages further validate and in strong agreement with several previous studies where various members of the heterotrophic bacterial populations, especially by the *Gammaproteobacteria and Alphaproteobacteria* classes were reported dominant within the coastal euphotic zones of marine environments (e.g., Wu et al. 2019). Follow-up investigations to this study would probably include the exploration of the coastal areas of the Indian Ocean that are also in close proximity to the cosmopolitan city of Cape Town as those of the Indian Ocean in order to compare the bacterial composition and diversity between these two oceans, especially at their points of confluence around the Cape of Agulhas.

## Acknowledgements

Sincere thanks to Mbulelo Mamani and the other staff of 15 on Orange, Autograph Collection Hotel in Cape Town for the support during field sampling at the Cape of Good Hope. Also, my appreciation goes to various staff members of CHUNLAB (Bioinformatics Made Easy, Seoul National University) for assistance during sequencing and bioinformatics data analysis. The study was generously supported by the Albion College Provost’s Office, Lori Duff and the award by the Hewlett-Mellon FDC funds made available during the sabbatical leave period that was utilized for this study.

